# Replication competent, 10-segmented influenza viruses as antiviral therapeutics

**DOI:** 10.1101/547059

**Authors:** Griffin D. Haas, Alfred T. Harding, Nicholas S. Heaton

**Author notes:** To whom correspondence should be addressed: Nicholas S. Heaton, PhD, Assistant Professor, Department of Molecular Genetics and Microbiology (MGM), Duke University Medical Center, 213 Research Drive, 426 CARL Building, Box 3054, Durham, NC 27710. Equal contribution (Griffin D. Haas, Alfred T. Harding).

## Abstract

Influenza A viruses (IAVs) encode their genome as eight negative sense RNA segments. During viral assembly, the failure to package all eight segments, or packaging of a mutated segment, renders the resultant virion incompletely infectious. It is known that the accumulation of these defective particles can limit viral disease by interfering with the spread of fully infectious particles. In order to harness this phenomenon therapeutically, we defined which viral packaging signals were amenable to duplication and developed a viral genetic platform which allowed the production of replication competent IAVs that package up to two additional artificial genome segments for a total of 10 segments. These artificial genome segments are capable of acting as “decoy” segments that, when packaged by wild-type (WT) viruses, lead to the production of non-infectious viral particles. Despite 10-segmented viruses being able to replicate and spread *in vivo*, these genomic modifications render the viruses avirulent. Excitingly, administration of 10-segmented viruses, both prophylactically and therapeutically, was able to rescue animals from normally lethally influenza virus infections. Thus, 10-segmented influenza viruses represent a potent anti-influenza biological therapy that targets the strain-independent process of viral assembly to slow the kinetics of productive viral spread and therefore limit viral disease.

**Author Summary:** Seasonal influenza infections are best prevented using vaccination. Vaccination, however, is not capable of completely preventing influenza infection, necessitating the use of anti-influenza therapeutics. To date, several different classes of anti-influenza therapeutics have been developed and used in order to combat these infections. Unfortunately, the incidence of influenza resistance to many of these therapeutics has begun to rise, necessitating the development of new strategies. One such strategy is to mimic the activity of naturally occurring viral particles that harbor defective genomes. These defective interfering particles have the ability to interfere with productive viral assembly, preventing the spread of influenza viruses across the respiratory tract. Furthermore, given the manner in which they target influenza segment packaging, a conserved feature of all influenza A viruses, resistance to this therapeutic strategy is unlikely. Here, we report the development of a genetic platform that allows the production of replication competent, 10-segmented influenza viruses. These viruses are capable of amplifying themselves in isolation, but co-infection with a wild-type virus leads to segment exchange and compromises the spread of both viruses. This interference, while mechanistically distinct from naturally occurring defective particles, was able to target the same viral process and rescue animals exposed to an otherwise lethal viral infection. This viral-based approach may represent a cost effective and scalable method to generate effective anti-influenza therapeutics when vaccines or anti-viral drugs become ineffective due to acquisition of viral resistance mutations.

## Introduction

Influenza virus infections represent a substantial global burden on human health. Each year, it is estimated that influenza viruses cause up to 5 million severe infections globally, resulting in up to 645,000 mortalities [1, 2]. In 2018, patient care and productivity loss due to influenza infection cost an estimated $11.2 billion in the U.S. alone [3]. Influenza A viruses (IAVs), the major contributor to total human influenza disease, possess a segmented genome consisting of eight discrete, negative-sense viral RNAs (vRNAs) [4]. Each of the eight vRNA segments consist of terminal 5’ and 3’ untranslated regions (UTRs) flanking an internal open-reading frame that encodes that one or maximally two viral proteins [5]. The UTRs, as well as the proximal portions of the coding regions, form “packaging signals” that are both necessary and sufficient for incorporation of each vRNA into progeny virions [6-11]. Although the underlying mechanisms that control packaging are incompletely understood, it has been hypothesized that segments may potentially interact with one another via vRNA-vRNA interactions across genome segments [12-16]. In any case, experimental evidence has supported the theory that non-random genome packaging controls the proper incorporation of segments into progeny virions [17, 18]. The segmented nature of the viral genome, and at least some intra-strain conserved regions of the packaging signals, allows for reassortment to occur between strains that have coinfected the same host cell [19]. The process of genetic reassortment, termed antigenic shift, can lead to the development of novel strains and can cause pandemic outbreaks, such as the one that occurred in 2009 with the H1N1 pandemic “swine” flu virus [20].

Currently, the primary measure used to control IAV spread is prophylactic immunization. However, due to rapid viral accumulation of point mutations, a process known as antigenic drift, vaccination can have limited efficacy. In these cases, healthcare providers must turn to therapeutic options for treating influenza disease. Adamantanes, matrix ion channel inhibitors, were the first IAV therapeutics developed, and were approved for clinical use in 1966 [21]. However, shortly after their deployment, it was apparent that IAVs were capable of developing rapid resistance to matrix ion channel inhibitors [22, 23]. High levels of resistance to adamantanes are now widespread in H1, H3, H5, H7, H9, and H17 subtype influenza A viruses, retiring the use of matrix ion channel inhibitors in treating influenza disease [22, 24]. Neuraminidase inhibitors, such as oseltamivir, are now the most commonly used IAV therapeutic [25]. As with adamantanes, this class of antiviral suffers from resistance as well [26-28]. In fact, greater than 90 percent of 2008-2009 pre-pandemic, globally circulating H1N1s were reported as having resistance to oseltamivir alone [21]. While these levels have decreased since the arrival of the 2009 pandemic-clade H1N1s, resistant strains are still isolated each year, highlighting the risk of widespread evolution of antiviral resistance [29, 30]. Finally, an mRNA cap snatching inhibitor, Baloxavir, was recently FDA-approved [31, 32], and the rate at which viral resistance may be acquired is currently unknown. Due to the incomplete efficacy of these therapeutics, as well as emerging viral resistance, additional antiviral therapeutics are in in various stages of development [31]. One promising alternative therapeutic approach, that theoretically would be difficult for IAVs to develop resistance to, has been the utilization of defective viral particles that disrupt viral replication and packaging [33].

Defective viral particles are not unique to influenza viruses, and research has demonstrated their formation and importance for a number of RNA viruses [34-36]. For influenza viruses, defective interfering particles, or DIPs, are replication-incompetent virions that frequently harbor one or more viral gene segments with a significant truncation of the open reading frame (ORF) of that segment [37]. Deletions can occur spontaneously during the replication stage of the viral lifecycle when the viral RNA-dependent RNA polymerase skips over a portion of the ORF, and generates a large deletion in that segment while still maintaining the 5’ and 3’ packaging signals necessary for gene segment incorporation [38]. If this partially deleted segment is packaged into nascent virions, virus particles are produced that are capable of infecting a host cell, but are then unable to produce subsequent viable progeny due to the lack of the protein normally encoded by the defective vRNA segment [39]. While DIPs are themselves replication incompetent, due to their defective segments, they can be successfully propagated during coinfection with a “helper” wild-type IAV. Although it was believed that such coinfections are relatively uncommon, recent work has shown that co-infection may actually help facilitate productive virus replication [40]. If DIP coinfection does occur, the defective segment(s) of the interfering particle are replicated more quickly than their wild-type counterparts due to their significantly smaller size [33, 41]. This rapid replication allows the defective segment(s) to outcompete the wild-type vRNAs for genome packaging, interfering with the ability of replication-competent wild type IAV progeny to be generated and spread.

The disruptive effect of DIPs has long garnered attention as a potential influenza antiviral [42-44]. Studies have shown that laboratory produced DIPs can be used prophylactically and therapeutically to protect mice from a lethal wild type IAV infection [45]. Further, coinfection of this same DI virus design with a dose of the 2009 H1N1 pandemic virus was found to reduce the symptoms of disease in a ferret model [46]. This DI system has also been shown to be effective *in vitro* in human respiratory tract cell lines [47]. Despite these advances, options for generating DIPs have been limited. Initially, DIPs were synthesized via high multiplicity passaging, which not only generates diverse DI populations with varying efficacy, but also contains wild-type IAVs that must also be inactivated by UV irradiation [48, 49]. Reverse genetic cloning has offered a means through which to generate populations of specific DIP genotypes, however this method requires the use of helper viruses for the proliferation of the DIPs, again necessitating a subsequent UV inactivation. Cell culture optimization for production of DIPs is under way, but is currently only able to produce high yield batches consisting of mixed DI populations with varying efficacy, or purer populations with a significantly reduced yield that are not sufficient for therapeutic use [50].

We were interested in generating IAV mutants that therapeutically mimic the inhibitory activity of DIPs, but using a fundamentally different molecular strategy. We hypothesized that the development of a live-attenuated virus harboring artificial genome segments could potentially be used to achieve this goal. This virus would be designed in such a way that the artificial segments would not interfere with its own replication, but when co-infection with a WT virus occurred, then cross-packaging of genome segments between the two viruses would lead to the production of non-viable particles and thus halt viral spread. In this report, we describe the generation of a live attenuated, 10-segmented (10S) influenza virus that accomplishes this goal. Administration of 10S viruses either prophylactically or therapeutically rescued animals from an otherwise lethal viral infection. Since the highly conserved, strain independent process of viral assembly is the target of this approach, we predict that it will be difficult or even impossible for IAV to generate functional escape variants through acquisition of random mutations.

## Results

### Evaluation of viral genetic manipulations capable of generating 9-segmented IAVs

We were interested in generating influenza viruses that could be genetically programmed to harbor artificial genomic segments that would interfere with the correct packaging of genome segments into nascent virions. It was previously reported that the NA packaging signals could be duplicated and utilized to package a ninth genomic segment [51]. While this approach was utilized to encode additional antigens as a vaccine platform, we theorized that this and similar approaches could be utilized to generate viruses harboring artificial, interfering segments. We therefore tested the ability to duplicate various packaging signals and generate 9-segmented viruses. We tested: NA, NP, HA, and PA duplicated packaging signals harboring different viral proteins in different combinations (Table 1). In all cases, the “9^th^” segment (Supplementary Figure 1) was designed with unique packaging signals so that it would always be packaged, but failure to package the duplicated packaging signal segment would lead to the loss of an essential viral protein. The 9^th^ segment always encoded super-folder GFP (sfGFP) or mCherry. Surprisingly, very few segment duplications were amenable to this approach. As previously reported, duplication of the NA packaging signal is tolerated, but of everything else tested, only duplication of the PA packaging signal was tolerated (Table 1).

**Table 1.** 9 segmented virus design strategies. A description of the manipulated packaging signals, encoded proteins, and success of rescuing each 9 segmented IAV strategy.

|  | Segment Design | Protein | Successful Rescue? |  | Segment Design | Protein | Successful Rescue? |
| --- | --- | --- | --- | --- | --- | --- | --- |
|  | **9s PB1-mCherry-PB1 / NA-PB1-NA Design** |  |  |  | **9s PB2-sfGFP-PB2 / PA-PB2-PA Design** |  |  |
| 1 | WT PB2 | PB2 | **Yes**<br>(Gao et al.) | 1 | **PA-PB2-PA** | PB2 | **Yes** |
| 2 | **NA-PB1-NA** | PB1 |  | 2 | WT PB1 | PB1 |  |
| 3 | WT PA | PA |  | 3 | WT PA | PA |  |
| 4 | WT HA | HA |  | 4 | WT HA | HA |  |
| 5 | WT NP | NP |  | 5 | WT NP | NP |  |
| 6 | WT NA | NA |  | 6 | WT NA | NA |  |
| 7 | WT M | M1, M2 |  | 7 | WT M | M1, M2 |  |
| 8 | WT NS | NS1, NEP |  | 8 | WT NS | NS1, NEP |  |
| 9 | **PB1-mCherry-PB1** | mCherry |  | 9 | **PB2-sfGFP-PB2** | sfGFP |  |
|  | **9s PB2-sfGFP-PB2 / NP-PB2-NP Design** |  |  |  | **9s M-zsGreen\|M2-M / HA-M1-HA** |  |  |
| 1 | **NP-PB2-NP** | PB2 | **No** | 1 | WT PB2 | PB2 | **No** |
| 2 | WT PB1 | PB1 |  | 2 | WT PB1 | PB1 |  |
| 3 | WT PA | PA |  | 3 | WT PA | PA |  |
| 4 | WT HA | HA |  | 4 | WT HA | HA |  |
| 5 | WT NP | NP |  | 5 | WT NP | NP |  |
| 6 | WT NA | NA |  | 6 | WT NA | NA |  |
| 7 | WT M | M1, M2 |  | 7 | **HA-M1-HA** | M1 |  |
| 8 | WT NS | NS1, NEP |  | 8 | WT NS | NS1, NEP |  |
| 9 | **PB2-sfGFP-PB2** | sfGFP |  | 9 | **M-zsGreen \| M2 - M** | zsGreen, M2 |  |
|  | **9S HA-sfGFP-HA / NS-HA-NS Design** |  |  |  | **9s NS-mCherry\|NEP-NS / NA-NS1-NA** |  |  |
| 1 | WT PB2 | PB2 | **No** | 1 | WT PB2 | PB2 | **Yes**<br>(Unstable) |
| 2 | WT PB1 | PB1 |  | 2 | WT PB1 | PB1 |  |
| 3 | WT PA | PA |  | 3 | WT PA | PA |  |
| 4 | **NS-HA-NS** | HA |  | 4 | WT HA | HA |  |
| 5 | WT NP | NP |  | 5 | WT NP | NP |  |
| 6 | WT NA | NA |  | 6 | WT NA | NA |  |
| 7 | WT M | M1, M2 |  | 7 | WT M | M1, M2 |  |
| 8 | WT NS | NS1, NEP |  | 8 | **NA-NS1-NA** | NS1 |  |
| 9 | **HA-sfGFP-HA** | sfGFP |  | 9 | **NS- mCherry \| NEP - NS** | mCherry, NEP |  |

Additionally, we also tested the potential of using splice sites in the 7th and 8th segments of IAV to generate 9S viruses expressing either M1 or NS1 in the ninth segment and a fluorescent protein with M2 or NEP in segment 7 or 8, respectively (Table 1, Supplementary Figure 1). These segments were designed so that viral splicing would remain intact. Duplicating HA packaging signals and encoding M1 as an artificial segment was unsuccessful, however duplicating the NA packaging signals and encoding NS1 successfully yielded 9S virions. The “NS” segment encoding mCherry and NEP however, immediately lost mCherry signal upon viral rescue, indicating that this approach is not useful for the stable incorporation of protein or nucleic acid. From these experiments we conclude that the duplication of packaging signals is not a generalizable approach for all segments, but in some specific cases, such as the with NA and PA packaging signals, this approach can be utilized to force viral packaging of a 9^th^ genomic segment.

### Characterization of 9-segmented viruses and their therapeutic efficacy

After determining which combinations of packaging signal duplications were tolerable, we began *in vitro* characterizations of the 9S PB1 mCherry virus, with duplicated NA packaging signals, and the 9S PB2 sfGFP virus, with duplicated PA packaging signals (Figure 1A & 1B). Multicycle growth experiments, both in embryonated chicken eggs and in MDCK cells, demonstrated that while both of these viruses exhibit attenuated levels of growth (Figure 1C), they do successfully package and propagate the artificial segment (Figure 1D). We hypothesized that the decreased viral growth may be due to the viruses only packaging one of the segments that harbors the duplicated packaging signal. If this were the case, we would expect to observe a large number of defective, 8 segmented viruses lacking an essential viral protein. In order to test this, we grew the viruses in embryonated chicken eggs and measured infectious particles via plaque assay, and we also performed a hemagglutinin (HA) assay, which measures both infectious and noninfectious particles (Figure 1E,F). Again, we observed a dramatic reduction in viral titer, however the magnitude of the observed defect in the HA assay was much smaller. To represent this difference, we calculated the “Relative DI units” of our 9S viruses, relative to WT virus, with WT set at an arbitrary value of 1, by dividing HA units by the endpoint titer (Figure 1G). As expected, the 9S PB1 mCherry virus produced ∼10^2^ times more non-viable progeny than did the WT PR8 virus, while the 9S PB2 sfGFP virus produced ∼10^3^ times more non-viable progeny than did WT PR8 (Figure 1G). Thus, both 9S viruses produced a significantly higher ratio of non-viable to viable virions than WT PR8 virus, and that ratio was, to some extent, dictated by which viral segment had been duplicated.

**Figure 1.**
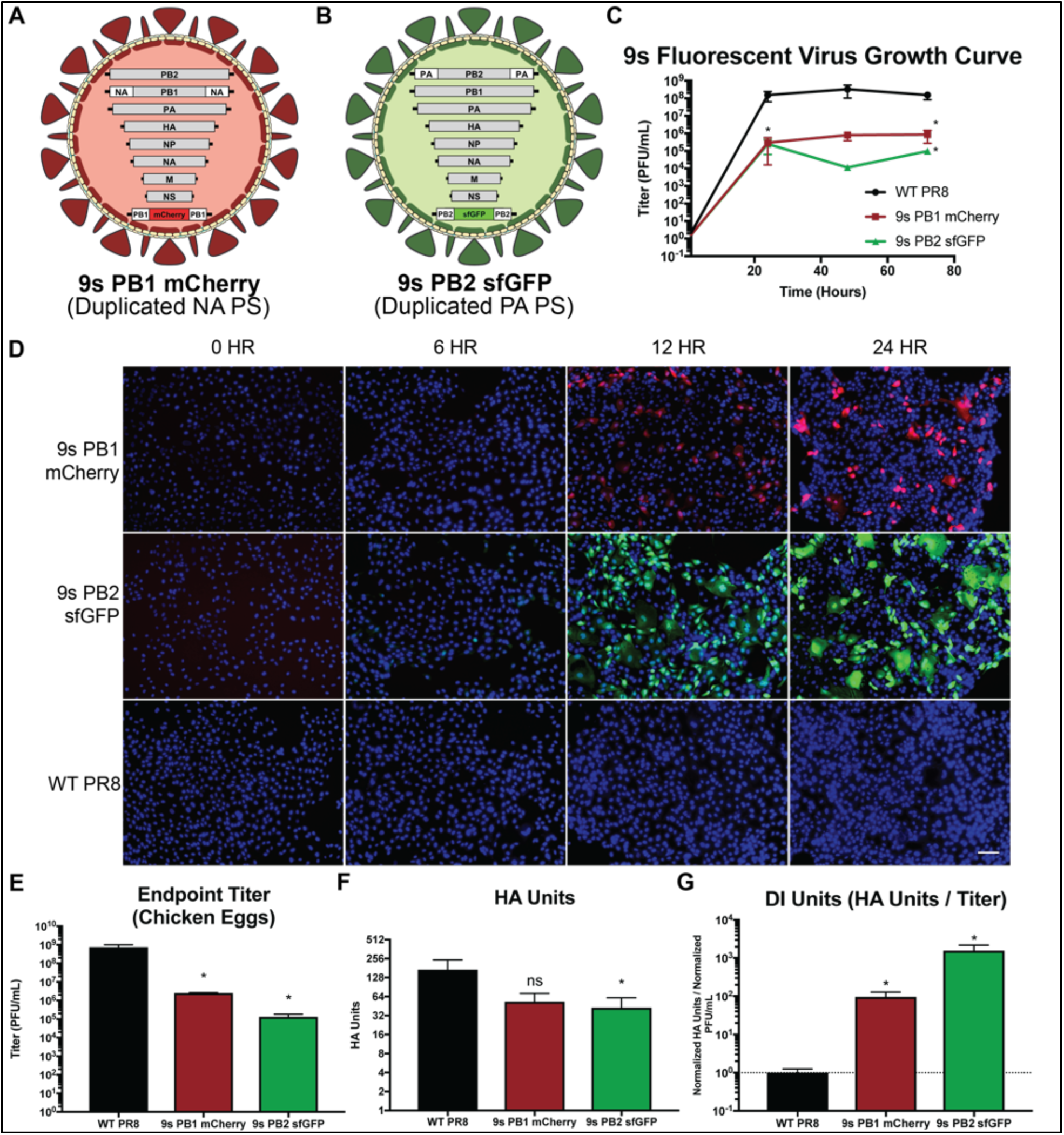
9-Segmented fluorescent viruses generate proportionally more defective interfering particles than WT IAV. (A and B) Genome design of the 9s PB1 mCherry virus (A) and the 9s PB2 sfGFP virus (B). (C) Growth curve of 9s PB1 mCherry (▄), 9s PB2 sfGFP (Δ), and WT PR8 (•) viruses titered in MDCK cells 0, 24, 48, and 72 hours post-infection in 10-day old embryonated chicken eggs. (D) Fluorescent microscopy images of 9s PB1 mCherry, 9s PB2 sfGFP, or WT PR8 virus-infected MDCK cells at 0, 6, 12, and 24 hours post-infection; nuclei were stained blue using DAPI staining and the scale bar represents 100 micrometers. (E) Endpoint titer 72 hours post-infection in 10-day old embryonated chicken eggs of the 9-segmented fluorescent viruses as compared to WT PR8 virus. (F) HA assay 72 hours post infection in 10-day old embryonated chicken eggs of the 9-segmented fluorescent viruses as compared to WT PR8 virus. (G) The “DI Units” of the 9-segmented fluorescent viruses as compared to that of WT PR8 virus, calculated by dividing respective normalized HA units by normalized endpoint titer. For all graphs, * represents a p-value of ≤ .05 and ** represents a p-value of ≤ .001.

We next wanted to assess how a 9^th^ segment affected the virulence of the virus, as well as assay the ability of the two 9S fluorescent viruses to modulate influenza disease. To determine if the addition of the 9^th^ segment attenuated the virus, LD50 experiments were performed in immunocompetent C57BL/6 mice. Wild-type PR8 virus was found to be lethal at all doses tested, killing all infected mice with as little as 10 PFU (Figures 2A,D). The two 9S viruses, however, were significantly attenuated relative to the parental virus. The 9S PB1 mCherry virus required 10^4^ PFU for lethality and 10^2^ PFU treated animals showed no death or weight loss (Figure 2B,E). Similarly, the 9S PB2 sfGFP caused lethal disease at a dose of 10^4^ PFU and 10^2^ PFU treated animals exhibited no death or weight loss (Figure 2C,F). The attenuation of the 9S viruses suggested that these viral genomic designs may fit the criteria of a live-attenuated therapeutic. We therefore assessed the capability of each 9-segmented fluorescent virus to interfere with a lethal challenge of WT PR8. For this initial test, we coinfected animals with 20 PFU of WT virus in combination with 500 PFU of either the 9S PB1 mCherry or the 9S PB2 sfGFP virus and monitored animals for body weight loss for 14 days post-infection (Figure 2G). 500 PFU of the 9S viruses was chosen as the highest dose that would not be expected to induce any clinical disease. Non-treated control animals rapidly lost body weight and succumbed to the challenge, as expected (Figure 2H, 2K). Administration of the 9S PB1 mCherry virus caused a measurable protective effect, with treated animals experiencing an ∼48 hour delay in the onset of body weight loss when compared to the lethal WT PR8 challenge (Figure 2I). Moreover, 25% of coinfected animals survived and recovered from this normally lethal challenge with WT PR8 (Figure 2L). In contrast, the 9S PB2 sfGFP virus did not cause treated animals to display any statistically significant reduction in weight loss or increased survival compared to the lethal WT PR8 challenge alone (Figure 2J, 2M).

**Figure 2.**
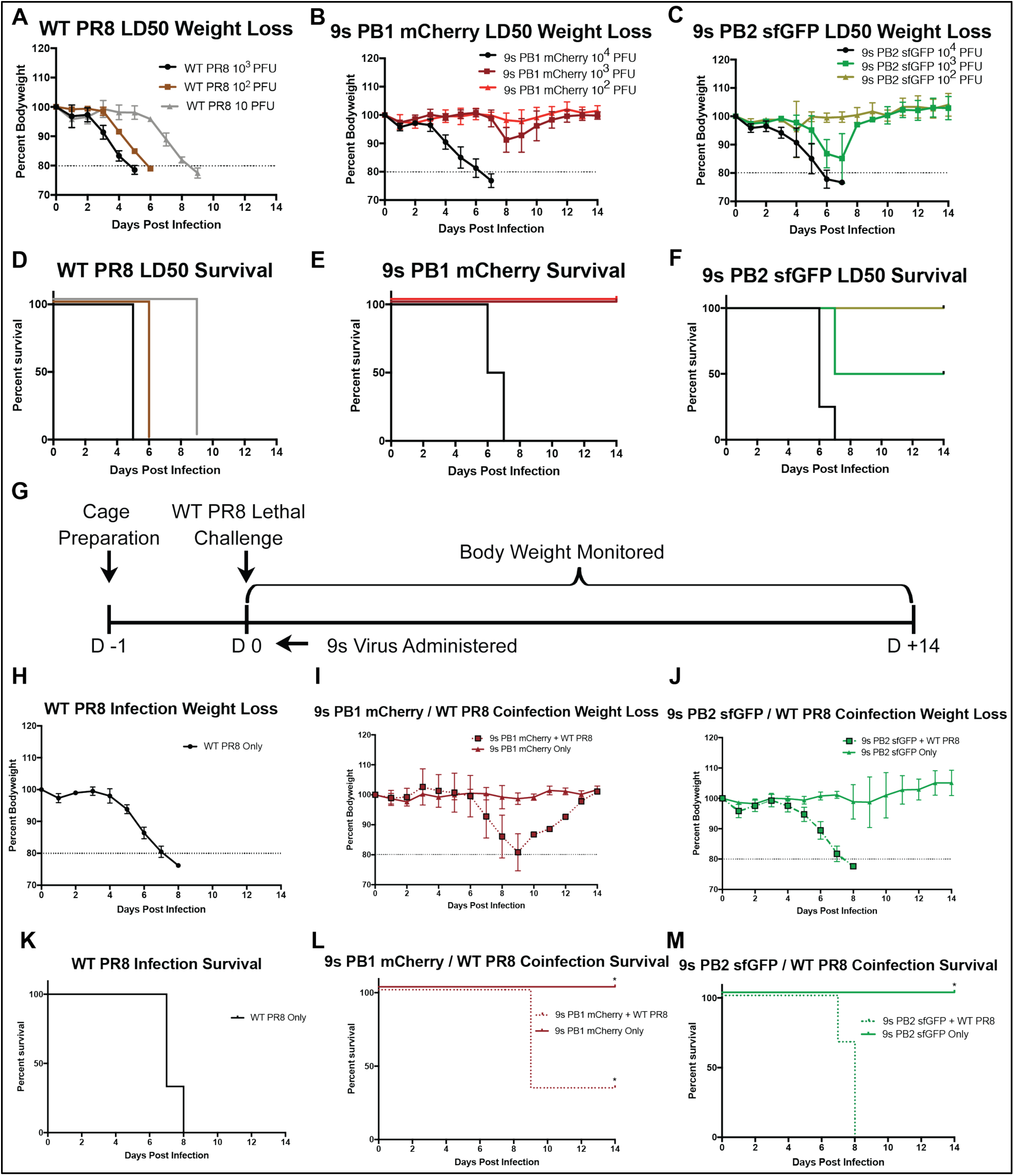
9-Segmented influenza viruses are highly attenuated and their administration at the time of infection can protect from lethal viral challenge. (A – C) Weight loss curves from infections with the indicated doses of WT PR8 virus (A), the 9s PB1 mCherry virus (B), or the 9s PB2 sfGFP virus (C). (D-F) Survival curves from infections with the indicated doses of WT PR8 virus (D), the 9s PB1 mCherry virus (E), or the 9s PB2 sfGFP virus (F). (G) Schema of C57BL/6J coinfection challenges. (H-J) Weight-loss curves from infecting mice with a lethal dose of WT PR8 ((•), 20 PFU) (H),a sublethal dose of the 9s PB1 mCherry virus ((▲), 500 PFU), or a lethal dose of WT PR8 virus in combination with 500 PFU 9s PB1 mCherry virus (?) (I),or a sublethal dose of the 9s PB2 sfGFP virus ((▲) 500 PFU) or a lethal dose of WT PR8 virus in combination with 500 PFU 9s PB2 sfGFP virus (?) (J). (K - M) Survival curves from coinfections challenging mice described in panels H - J, respectively.

### The artificial viral segment size is not correlated with therapeutic effect

While the 9S PB1 mCherry virus did offer some therapeutic effect, the effect size was minimal. We hypothesized that this was likely due to the design of our segment. During normal WT replication, DI segments arise from the large scale deletion of ORFs, often reducing the size of a DI segment to a total of less than 500 nucleotides. This significant reduction in size causes the DI segment to be replicated much faster than the full-length WT segment, drastically enhancing the chance that the DI segment is packaged into a progeny virion over the WT one. Our artificial segments, however, were actually larger than a standard DI segment, potentially reducing the efficacy of this strategy. In order to determine if the protective effect of a 9S virus could be augmented by making it more like a DIP, we designed a DI-like oligonucleotide to replace PB1 mCherry, based on a previously characterized PB1 DI segment reported by Saira *et al.* [38] (Figure 3A). We chose to focus on the PB1-mCherry segment as this segment showed a larger degree of protection from challenge relative to the PB2-sfGFP segment. We used the PB1 DI segment was in place of the mCherry expressing segment to generate a virus harboring a more DI-like segment (Figure 3B). The 9S PB1 DI virus again was attenuated relative to WT viruses by approximately the same magnitude as the other 9S viruses (Figure 3C). Analysis of titer and HA units after growth in chicken eggs revealed that similarly to the other 9S viruses, the 9S PB1 DI virus produced roughly 10^2^ times more non-viable progeny than WT PR8 virus (Figure 1D-F).

**Figure 3.**
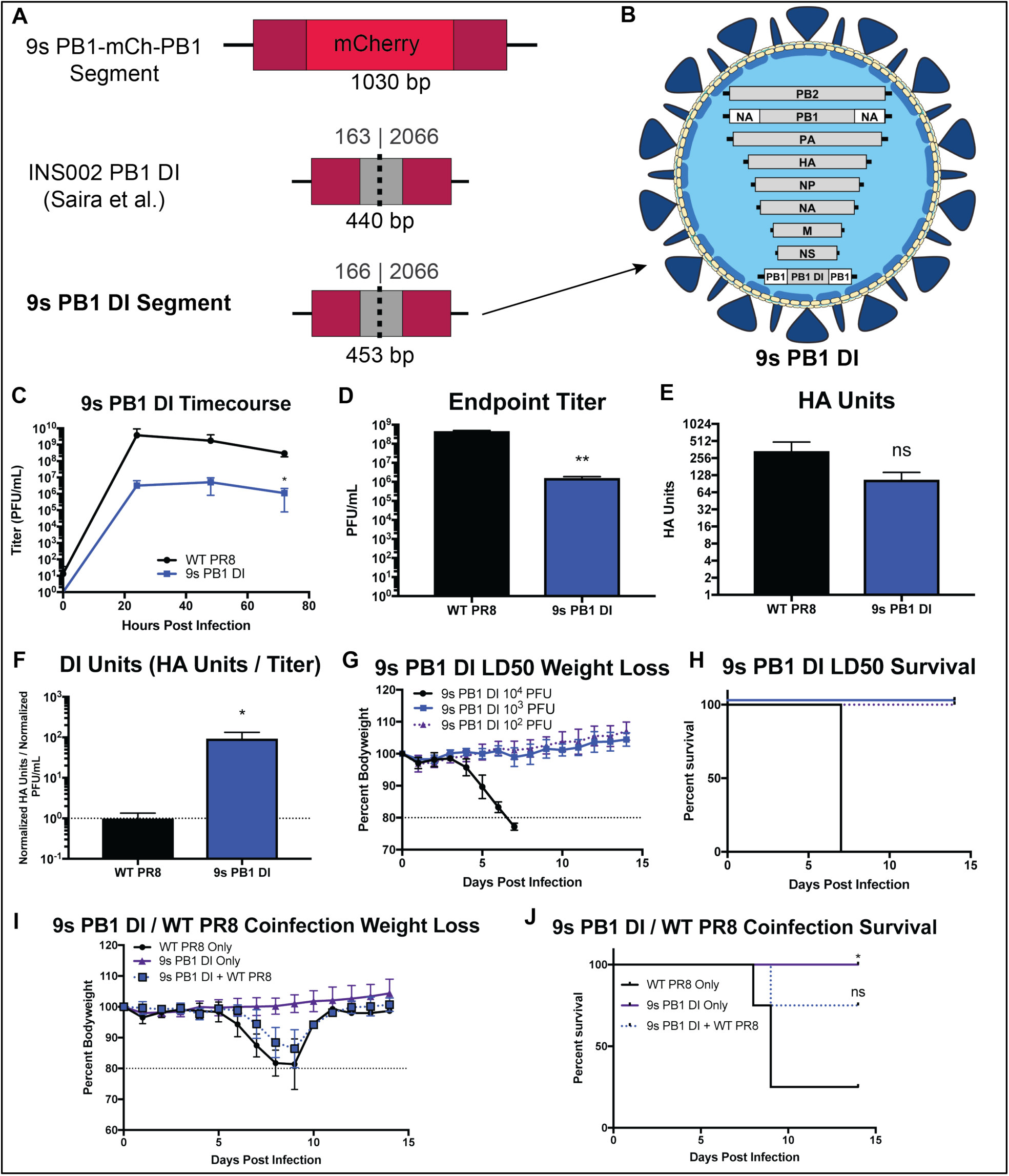
9-Segmented influenza viruses can harbor a natural defective interfering-like segment. (A) A schematic comparing the 9s PB1-mCherry-PB1 segment and the INS002 PB1 DI segment, which acted as a basis for the design of the 9s PB1 DI segment. (B) A schematic detailing the genome design of the 9s PB1 DI virus, including a ninth PB1-DI-PB1 segment. (C) Growth curve of the 9s PB1 DI virus titered in MDCK-cells at 0, 24, 48, and 72 hours post-infection in 10-day old embryonated chicken eggs. (D) Endpoint titer 72 hours post-infection in embryonated chicken eggs of the 9s PB1 DI virus as compared to WT PR8 virus. (E) HA assay 72 hours post infection in 10-day old embryonated chicken eggs of the 9s PB1 DI virus as compared to WT PR8 virus. (F) The “DI Units” of the 9s PB1 DI virus as compared to that of WT PR8 virus, calculated by dividing normalized HA units by normalized endpoint titer. (G) Weight loss curves from infection with the indicated doses of the 9s PB1 DI virus. (H) Survival curves from infections with the indicated doses of 9s PB1 DI virus. (I) Weight-loss curves from infecting mice with a sublethal dose of the 9s PB1 DI virus ((▲), 500 PFU), a lethal dose of WT PR8 ((•), 20 PFU), or a lethal dose of WT PR8 virus in combination with 500 PFU 9s PB1 DI virus (?). (J) Survival curves from infections described in panel I. For all graphs, * represents a p-value of ≤ .05 and ** represents a p-value of ≤ .001.

As expected, the 9S PB1 DI virus was significantly attenuated *in vivo*, even more so than the previous 9S viruses. Only the highest dose tested, 10^4^ PFU, was lethal, whereas the other two doses, 10^3^ and 10^2^ PFU, caused no death or weight loss (Figure 3G-H). To test the protective efficacy of the 9S DI virus, we simultaneously treated mice with 500 PFU of the DI virus together with a normally lethal dose of WT PR8. Similar to the 9S PB1 mCherry virus, the 9S PB1 DI virus was found to confer a protective effect, with weight loss occurring 24 hours later than seen in the control, WT PR8 challenged mice, and an increase in survival rates (Figure 3I,J). Thus, the 9S PB1 DI virus had a very similar protective effect to the 9S PB1 mCherry virus, suggesting that the ability of these viruses to interfere with influenza disease is independent of the artificial genome segment size.

### 10-segmented IAVs are viable and their administration can rescue infected animals from lethal viral disease

Since the size of the DI segment did not appear to play a critical role in interfering with viral replication/packaging, we hypothesized that potentially increasing the number of segments would increase the ability of the virus to interfere with WT viral spread. Given that we able to successfully utilize two 9S genome packaging strategies, that utilized distinct packaging signal duplications, to generate two different viable 9S IAV variants, we considered the possibility of combining the two to generate a viable 10S IAV. Indeed, these two strategies were compatible, and we were successful in rescuing a 10S IAV harboring 6 WT segments alongside four genetically manipulated ones (Figure 4A). Growth curve analysis of the 10S virus shows it is extremely attenuated, even more than the 9S viruses (Figure 4B). A fluorescent microscopy time course of the 10S virus revealed that virus indeed functionally co-packaged both of the fluorescent artificial segments at all timepoints (Figure 4C). Analysis of viral titer and HA units after growth in chicken eggs demonstrated that the 10S virus, while highly attenuated, produced a significantly higher ratio of nonviable progeny, nearly 10^3^ times higher than WT PR8 virus (Figure 4D-F).

**Figure 4.**
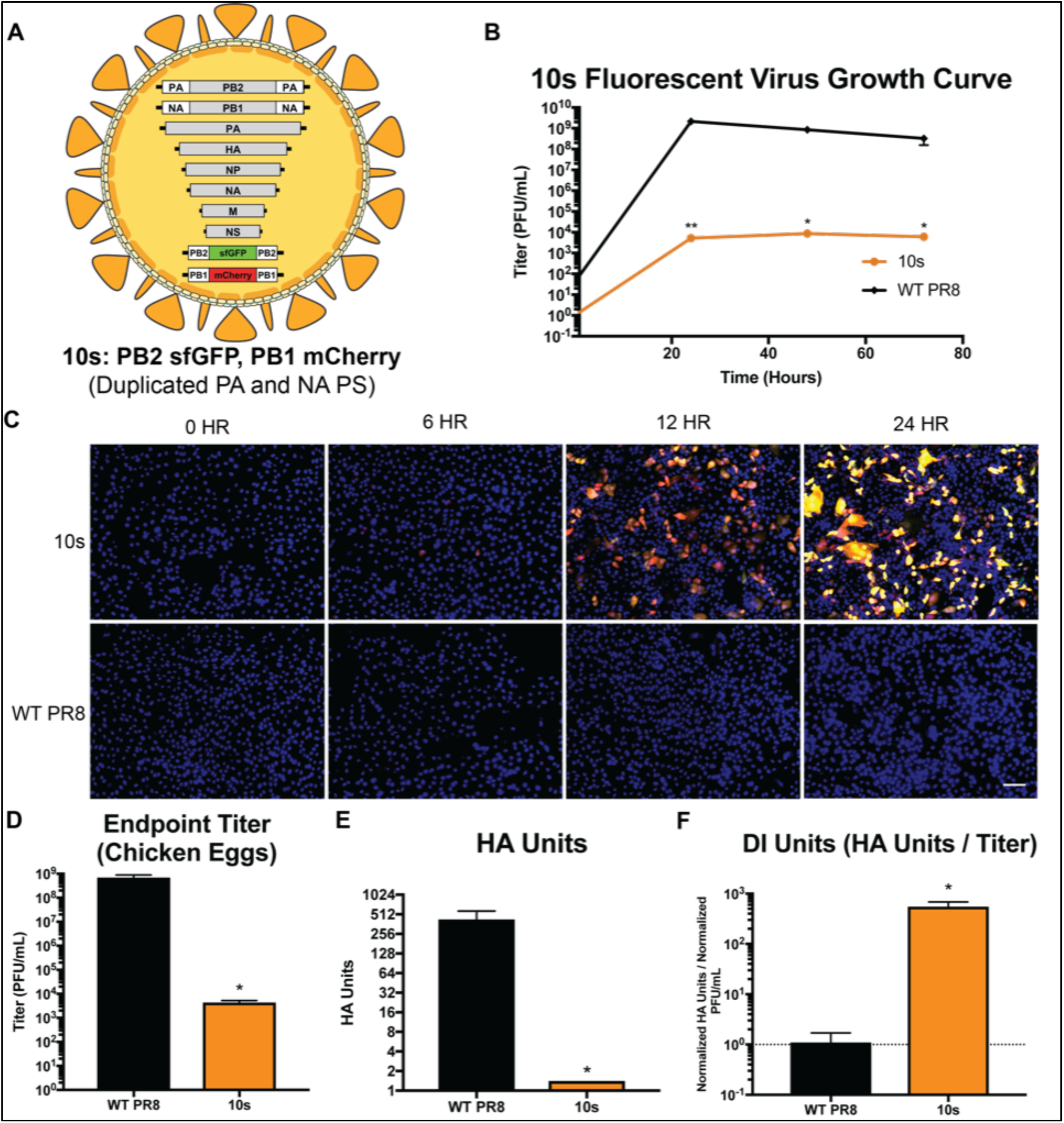
10-Segmented fluorescent viruses can be generated by combining two 9-segmented approaches. (A) Genome design of the 10s PB2 sfGFP PB1 mCherry virus. (B) Growth curve of the 10s virus measuring titered in MDCK cells at 0, 24, 48, and 72 hours post-infection in 10-day old embryonated chicken eggs as compared to PR8 WT (C) Fluorescent microscopy images of 10s or WT PR8 virus-infected MDCK cells at 0, 6, 12, and 24 hours post-infection; nuclei were stained blue using DAPI staining, and the scale bar represents 100 micrometers. (D) Endpoint titer 72 hours post-infection in 10-day old embryonated chicken eggs of the 10s virus as compared to WT PR8 virus. (E) HA assay of the 10s virus as compared to WT PR8 virus. (F) The “DI Units” of the 10s virus as compared to that of WT PR8 virus, calculated by dividing normalized HA units by normalized endpoint titer. For all graphs, * represents a p-value of ≤ .05 and ** represents a p-value of ≤ .001.

Next, we assessed the virulence of the 10S virus via an LD50 experiment in C57BL/6 mice. The virus was highly attenuated; animals infected with doses as high as 10^4^ PFU, which caused mortality when using any of the 9S viruses, experienced no detectable decline in body weight and all survived (Figure 5A-B). This increased attenuation is highly desirable when considering its use as a potential antiviral therapeutic. We were concerned, however, that this attenuated replication level would be too low to demonstrate any protective efficacy against WT IAV. As an initial test of the potential efficacy of a 10S virus as an antiviral agent, we challenged C57BL/6 mice with 20 PFU of WT PR8 virus in combination with 5000 PFU of the 10S virus (Figure 5C). Remarkably, animals infected with both WT PR8 virus and 10S virus exhibited no detectable weight loss, whereas WT PR8 only infected control animals began to lose body weight as early as 5 days post-infection (Figure 5D). All of the 10S treated animals survived the infection, whereas all of the WT PR8 only infected animals succumbed (Figure 5E). We were next curious to assess 10S virus efficacy in more authentic therapeutic application. We therefore infected mice with a lethal dose of 20 PFU of WT PR8 virus, administered the 10S therapeutic dose 24 hours later, and then monitored animals for weight loss for 14 days post-infection (Figure 5F). While the 10S virus treatment 24 hours after WT infection was not as effective as a simultaneous coinfection, we did observe a significant reduction in weight loss of the 10S treated animals, when compared to animals infected with WT PR8 alone (Figure 5G). Furthermore, animals that were administered the 10S virus at 24 hours after WT PR8 infection had a significantly increased survival rate, 25% mortality versus 100% mortality, when compared to control WT PR8 only infected animals (Figure 5H). Thus, we have developed an approach to generate viable 10S viruses and have shown that administration of 10S viruses either at the time of infection with WT IAV, or up to 24h later, can effectively prevent, lethal influenza virus disease.

**Figure 5.**
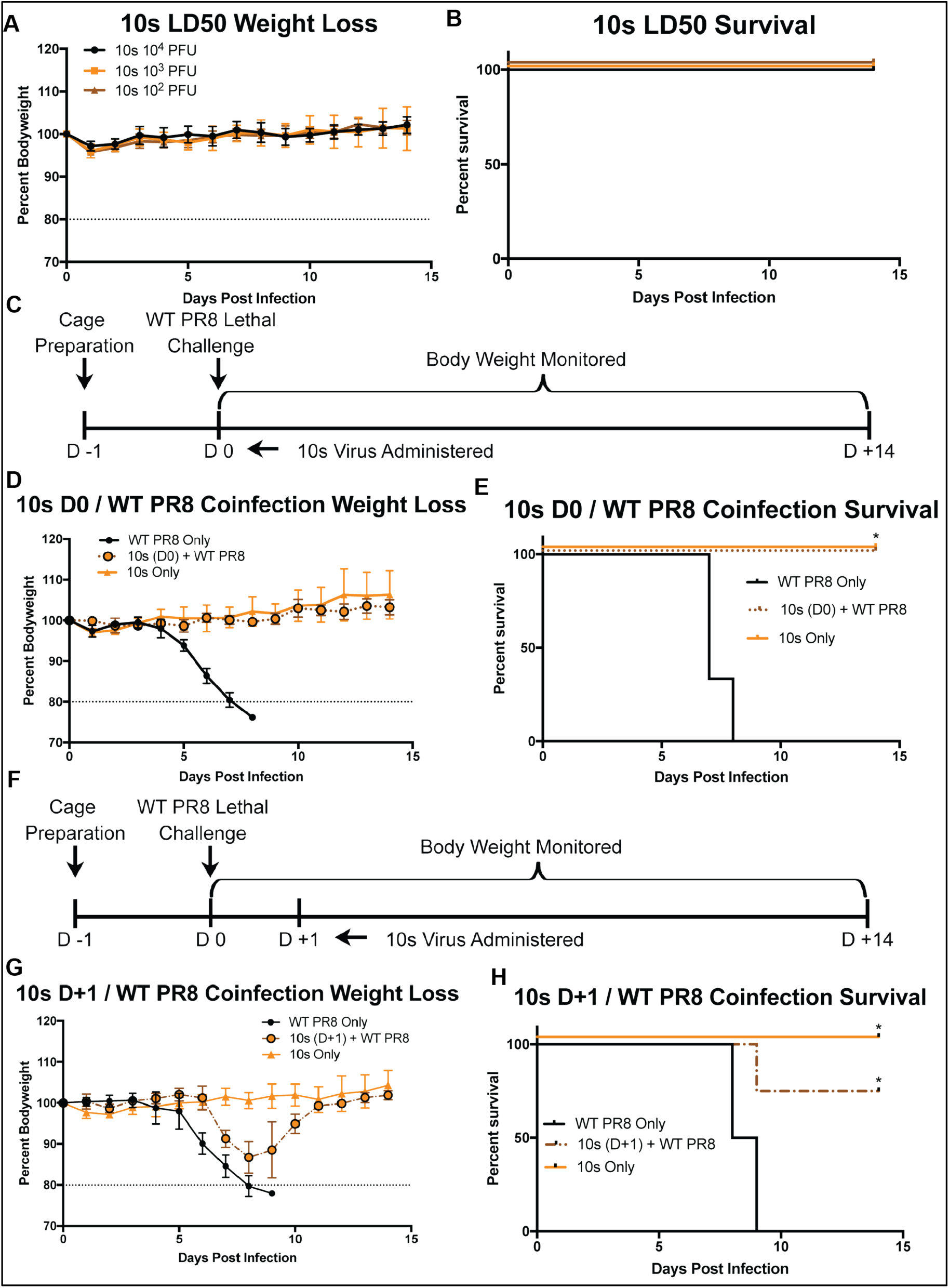
10-Segmented influenza viruses are highly attenuated and protect from lethal viral challenge when administered therapeutically. (A) Weight loss curves from infections with the indicated doses of 10s virus. (B) Survival curves from infections with the indicated doses of 10s virus. (C) Schema of C57BL/6J coinfection challenge at D0. (D) Weight loss curves from infecting mice with a sublethal dose of the 10s virus ((▴), 5000 PFU), a lethal dose of WT PR8 ((•), 20 PFU), or a lethal dose of WT PR8 virus in combination with 5000 PFU 10s virus (?). (E) Survival curves from the infection groups described in panel D. (F) Schema of C57BL/6J therapeutic 10s administration at 24 hours post-infection with lethal dose of WT PR8 virus. (G) Weight loss curves from infecting mice with a sublethal dose of the 10s virus ((▲), 5000 PFU), a lethal dose of WT PR8 ((•), 20 PFU), or a lethal dose of WT PR8 virus in combination with a dose of 5000 PFU 10s virus administered 24 hours later (?). (H) Survival curves from the infection groups described in panel G.

**Figure 6.**
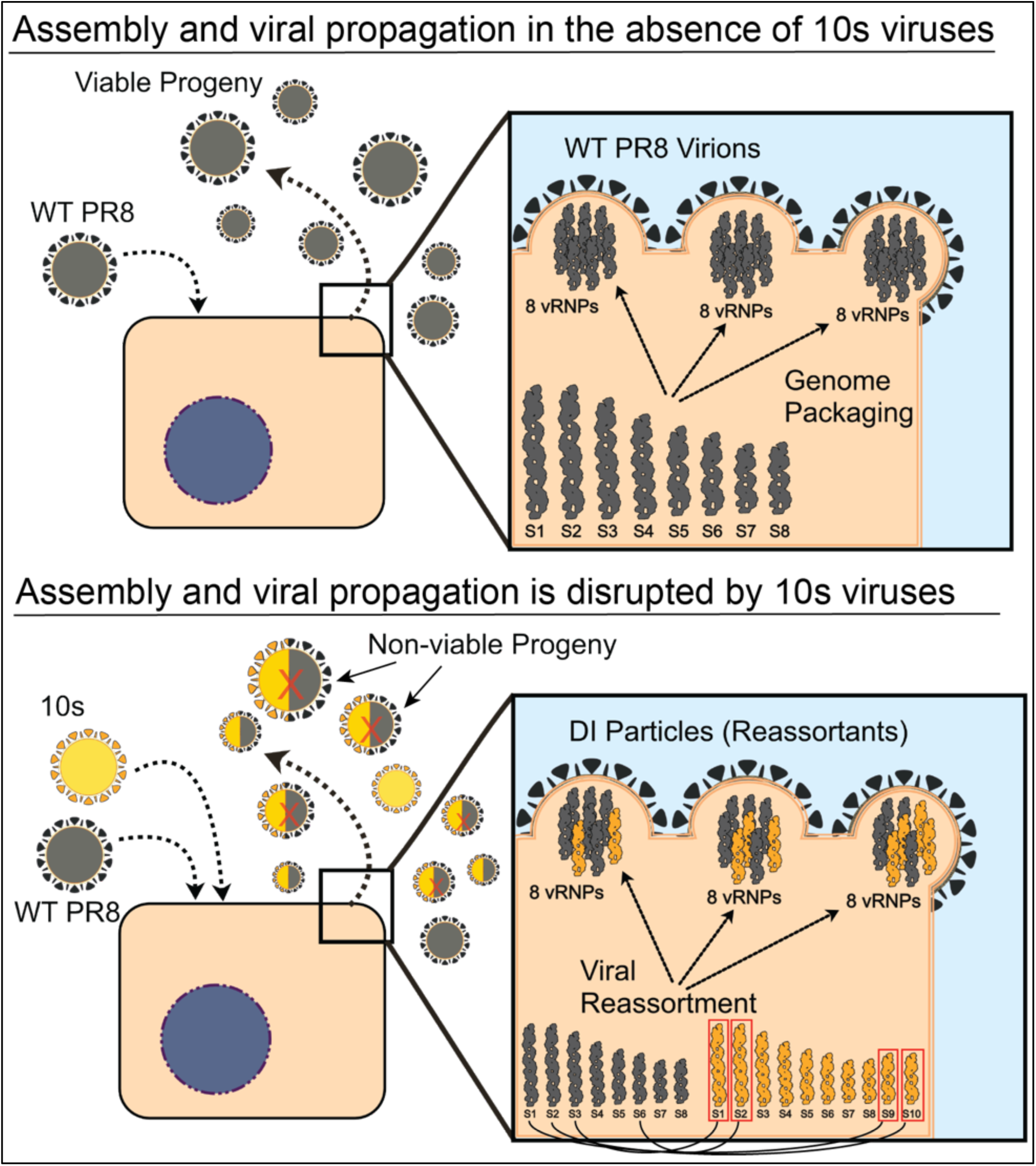
Model for 10s viral interference with WT viral spread. WT PR8 virus (grey) replication produces viable progeny (top panel). 10s virus (orange) coinfection with WT PR8 virus facilitates incomplete genome packaging, resulting in disrupted WT PR8 replication and the production of non-viable progeny (lower panel). The black curved lines indicate packaging signal equivalence between the WT and 10s virus genomic segments, and incorporation of any of the four red-boxed segments into WT virions will generate non-viable genomic reassortants.

## Discussion

This research was initially started with the goal of creating a replication competent, live attenuated virus that would be able to encode genomic segments capable of disrupting effective genomic packaging of a co-infecting WT virus. Our approach is mechanistically distinct from naturally occurring DI particles, which are naturally generated via large deletions of a viral segment. The 10S platform however, essentially mimics the concept of facilitating packaging of a defective viral segment, which then leads to the release of virions bearing incomplete viral genomes. In order to produce this virus, we first verified that a previously published approach of duplicating the NA segment packaging signals could be utilized to make the virus encode a 9^th^ genomic segment [51]. We next expanded upon that work and tested a variety of other genomic organizations and found that only rare combinations of viral genes and packaging signals were able to be tolerated by the virus. There are probably a number of constraints that underlie this phenomenon. First there is known to be a hierarchy of viral segment packaging [9], and thus, some segments (distinguished by the virus based on the packaging signals) may be less tolerant of duplication than others and lead to a disruption of the structure/assembly of the IAV genome. In line with this concept, work using 7-segmented influenza viruses has demonstrated that the requirement for different packaging signals is variable with respect to viral assembly [52]. Interestingly, this earlier work demonstrated that both the NA and PA packaging signals are not required for the packaging of the other genomic segments. Our ability to duplicate both of those packaging signals agrees with the concept that these particular packaging signals play a relatively less important role in viral assembly.

The 7-segmented virus work however, does not necessarily predict the ability of a given packaging signal to be duplicated. For example, NS packaging signals were also shown to be dispensable, yet we were unable to rescue a virus with duplicated NS packaging signals (Table 1). This discrepancy may be explained by the fact that the levels of transcription and translation of these viral segments is controlled by motifs in these specific segments [53, 54], and thus the combination of different viral ORFs and packaging signals leads to a disruption of the normal controllers of viral transcription/translation rates, negatively impacting viral fitness. This concept is somewhat supported by our data that a virus encoding the PB2 protein flanked by NP packaging signals is non-viable; NP is expressed in cells to a much higher level than PB2. When we encoded PB2 flanked by PA packaging signals however, the virus was viable, and PA and PB2 levels in the infected cell are reasonably similar [55].

To our knowledge, 9S viruses had never previously been tested for their ability to interfere with IAV disease progression, and we therefore decided to test our 9S viruses in that capacity. We chose to administer these viruses at the time of infection with a lethal dose of WT virus as a reasonably stringent test for potential efficacy of the approach. Disappointingly, only one of our 9S viruses displayed any protective efficacy, and the effect was limited. In order to try and improve the ability of our artificial viral segments to titrate viral RdRPs away from WT genomic segments, we made the artificial segment much smaller. Naturally occurring defective interfering viral segments are much smaller than our fluorescent protein encoding artificial 9^th^ segments, and we therefore generated a 9S virus harboring a segment that was more similar in size to naturally occurring DI segments [38]. While we were able to generate viruses that harbored these DI like segments, we found that the reduction in segment size led to very little, if any, improvement in efficacy. Although these insights are derived from a highly artificial system, our data suggest that it may be interesting to reevaluate the relative importance of segment length in the context of naturally occurring DI particles.

Since varying the artificial segment size was not correlated with protection from IAV, we hypothesized that the efficacy of our approach was instead dependent on the efficiency of packaging of our artificial genome segments by WT viruses, leading to progeny virions with incomplete genomes. Were that the case, making a virus which encoded more artificial segments could potentially confer higher protective efficacy. We therefore produced a 10S IAV that possessed 2 artificial genome segments instead of 1. Since our two validated genetic approaches were compatible with each other (i.e. different segments and packaging signals were utilized in the two approaches), we attempted to combine our 9s PB1 mCherry and 9s PB2 sfGFP virus strategies to produce a 10S virus.

This effort was successful, resulting in the first known report of a stable 10S IAV. Although the growth rates of 10S viruses were significantly reduced relative to both WT and 9S viruses, the protective effect observed was far superior to that seen with any of the 9S viruses. When administered at the same time of infection, 100% of our treated mice survived a normally lethal dose of WT PR8 while exhibiting no detectable weight loss. While the effect of truly therapeutic administration 24 HPI had a less striking effect, we were still able to significantly delay the onset of clinical symptoms and reduce mortality rates by up to 75%.

Aside from the therapeutic potential of the 10s virus, its generation raises a multitude of interesting questions that warrant additional study. Perhaps most obvious is the question of IAV genome architecture. It is well accepted that IAVs package their segments in a “pinwheel” or 7+1 conformation, wherein a single segment, most likely one of the polymerase segments based on its size, is packaged in the center with the remaining 7 segments arranged around it in a circular shape [56]. The genomic architecture of both 9S and 10S IAVs, however, has not yet been evaluated. Understanding how the addition of one, or even two, segments impacts this structure could lead to a much better comprehension of both its assembly and stability during IAV packaging. Along these same lines, it has been shown that these genomic segments are tightly organized within the viral particle, leaving little room for excess genomic material [57]. The ability to generate 9S viruses, let alone 10S, viruses raises important questions as to the maximum amount of genetic material IAV virions can hold. This becomes an especially important question when considering the potential for utilizing influenza viruses as viral vectors, a platform that has been used for delivering a wide variety of proteins and nucleic acids [58].

Finally, the fact that 9S and 10S viruses can interfere with WT viral propagation strongly supports the notion that cellular co-infection is a common occurrence *in vivo*. Despite the historical notion that most viral particles are fully infectious, likely due to the fact that IAV particles package all eight genomic segments the majority of the time [59, 60], recent work has suggested that co-infection may actually be not only a frequent occurrence that allows viral reassortment [61], but also a critical aspect of normal viral spread across infected tissues [62]. Since our 9S or 10S interfering effects are dependent on co-infection with WT viruses, we not only favor this model, but propose that even distinct viral infections that begin at different times are also subject to this co-infection phenomenon.

In summary, we have successfully defined genomic architectures that allow influenza viruses to harbor up to two additional, artificial segments. Our work suggests that not all IAV packaging signals are amenable to manipulations such as duplication, and that the particular characteristics of a “defective” viral segment are not as important to its interfering effect as the absolute number of segments that can disrupt productive genomic packaging. Continued development of the 10-segmented replication-competent IAV platform may lead to a novel class of therapeutics that can be easily manufactured, safely administered, and display protective efficacy against viruses that have evolved resistance to other antiviral therapies.

## Acknowledgements

We would like to thank Dr. Bryan Cullen and Dr. Alanson Girton for helpful discussions and critical reading of this manuscript. We are also grateful for contributions made by Heather Froggatt (Duke University) and Calla Telzrow (Duke University) for early work on strategies to develop 9-segmented viruses. N.S.H. is partially supported by federal funds from the National Institute of Allergy and Infectious Diseases, National Institutes of Health, Department of Health and Human Services, under CEIRS Contract No. HHSN272201400005C. A.T.H was supported by NIH training grant T32-CA009111.

## Materials and Methods

### Ethics Statement

All procedures were carried out in compliance with the Duke University IACUC approved protocol number A189-18-08. Duke University maintains an animal program that is currently registered with the USDA, assured through the NIH/PHS, and accredited by AAALAC International. Animals were monitored daily for the following: respiratory rate, ambulating difficulty, ruffled fur, lack of grooming, restlessness, reluctance to move, and bodyweight loss. Humane endpoints were primarily based on bodyweight loss, and defined as ≥ 20% of the starting bodyweight. The primary euthanasia method used was CO_2_ asphyxiation, followed by bilateral thoracotomy as a secondary method. SPF embryonated chicken eggs were purchased from Charles River Labs and incubated in the laboratory for viral stock amplification. Eggs were injected with virus 8-10 days post-fertilization and incubated until a maxim age 13 days post-fertilization.

### Animal Infections

Eight to 10-week-old C57BL/6 mice were purchased from Jackson Laboratories and maintained at Duke University animal facilities. For all experiments, a sample size of at least 4 mice per group were used. Prior to infection, mice were anesthetized with a 100-microliter injection of a ketamine-xylazine mixture. Mice were weighed and tail-marked, and 40 microliters of virus diluted in pharmaceutical-grade PBS was administered intranasally. Mice were weighed daily and euthanized once the predetermined humane endpoint was reached. Therapeutic doses of 9S and 10S half were ½ of the highest dose that caused only mild disease in mice when administered alone.

### Cell Culture

Madin-Darby canine kidney (MDCK) cells, from the American Type Culture collection (ATCC), were grown in minimal essential medium (MEM) supplemented with 10% fetal bovine serum, HEPES, NaHCO3, GlutaMAX, and penicillin-streptomycin. Human embryonic kidney 293T cells (from the ATCC) were grown in Dulbecco’s modified Eagle’s medium (DMEM) supplemented with 10% fetal bovine serum, GlutaMAX, and penicillin-streptomycin. All cells were cultivated at 37°C, at a 5% CO_2_ content, in the humidity controlled Heracell™ VIOS 160i Thermo Scientific incubators.

### Cloning and rescue of 9s and 10s viruses

Recombinant influenza viruses were generated as previously described by use of the ambisense pDZ rescue plasmid system [63]. The 9s PB1 mCherry segment was generated similarly to the construct described in [52], replacing GFP with mCherry. Briefly, the mCherry fluorescent protein coding sequence, preceded by a Kozak sequence (gccacc), was cloned into a PR8 PB1 packaging vector using PCR and subsequent NEBuilder® HiFi DNA Assembly reaction. The PB1 packaging vector consisted of nucleotides 1-146 of the 5’-most PB1 sequence (with all ATG start sites mutated) followed by an EcoRV site. A PmeI restriction site separated the 3’-most PR8 PB1 sequence of nucleotides 2189-2341. The PB1 coding sequence (nucleotides 25-2298), with a 5’ Kozak sequence and was cloned into a PR8 NA packaging vector using PCR and a subsequent NEBuilder® HiFi DNA Assembly reaction. The PB1 coding sequence 5’-most 75 nt and 3’-most 81 nt were silently mutated to remove packaging signal activity. The NA packaging vector consisted of nucleotides 1-173 of the 5’-most PR8 NA sequence (with all ATG start sites mutated) followed by an EcoRV site. A PmeI sequence separated the subsequent 3’-most NA sequence of nucleotides 1205-1413. Packaging vectors and primers were synthesized as ordered through Integrated DNA Technologies, Inc.

The 9s PB2 sfGFP segment was generated as follows: The sfGFP fluorescent protein coding sequence, preceded by a 5’ Kozak sequence, was cloned into a PR8 PB2 packaging vector using PCR and a subsequent NEBuilder® HiFi DNA Assembly reaction. The PB2 packaging vector consisted of nucleotides 1-158 of the 5’-most PB2 sequence (with all ATG start sites mutated) followed by a NheI site. An XhoI sequence separated the 3’-most PB2 sequence of nucleotides 2189-2341. The PB2 coding sequence (nucleotides 25-2298) with a 5’ Kozak sequence was cloned into a PR8 PA packaging vector using PCR and a subsequent NEBuilder® HiFi DNA Assembly reaction. The PB1 coding sequence 5’-most 30 nt and 3’-most 85 nt were silently mutated to remove packaging signal activity. The PA packaging vector consisted of nucleotides 1-129 of the 5’-most PA sequence (with all ATG start sites mutated) followed by an EcoRV site. A PmeI sequence separated the subsequent 3’-most PA sequence of nucleotides 2050-2233.

The 9s PB1 DI segment is based upon characterization of INS002, as described in [38]. DNA was synthesized via Integrated DNA Technologies, Inc. in which the aforementioned PB1 packaging vector contained PR8 PB1 nucleotides 146-190 followed by nucleotides 2094-2188. In the 5’-most region of this construct, all ATG start codons were mutated to prevent undesired translation initiation.

9S viruses were generated by transfecting two artificial segment plasmids into low-passage 293T cells alongside the 7 additional wildtype segment plasmids. Transfections were conducted using 12 microliters of MIRUS Mirus Trans-IT LT1 reagent in 200 microliters of OPTI-MEM. Rescue supernatant was collected after 24 hours of incubation, with 200 microliters injected into 10-day-old chicken eggs purchased from Charles River Laboratories, Inc. Eggs were allowed to incubate virus for 72 hours prior to collection of allantoic fluid. Viruses were purified via plaquing in MDCK cells and subsequent amplification in chicken eggs.

The 10-segment PB2 sfGFP, PB1 mCherry virus was generated by transfecting the recombinant PB2-sfGFP-PB2, PA-PB2-PA, PB1-mCherry-PB1, and NA-PB1-NA plasmids into low-passage 293T cells alongside each of the necessary 6 additional wildtype segment plasmids (PA, HA, NP, NA, M, NS). Transfections were conducted using 14 microliters of Mirus Trans-IT LT1 reagent in 200 microliters of OPTI-MEM. Rescue supernatant was collected after 24 hours of incubation, with 200 microliters injected into 10-day-old chicken eggs. Eggs were allowed to incubate virus for 72 hours prior to collection of allantoic fluid. Viruses were purified via plaquing in MDCK cells and subsequent amplification in chicken eggs. Stocks of concentrated 10s virions were prepared using a 30% sucrose cushion for 1 h at 25,700 rpm on the Sorvall TH-641 swinging bucket rotor.

### Viral titering

Allantoic fluid was collected from chicken eggs following infection, and viral titer was determined via standard plaque assay procedures on MDCK cells. Briefly, cells were incubated for 1 h in 500 microliters of diluted virus suspension at 37°C, before removing the virus and applying the agar overlay. Cells were then incubated at 37°C for 72 h before being fixed in 4% paraformaldehyde (PFA) in phosphate-buffered saline (PBS) for at least 4 h. The 4% PFA was then aspirated, and the agar layer was removed before washing cells in PBS and incubating them at 4°C overnight in mouse serum from PR8-infected mice. Mouse serum was diluted 1:2,000 in antibody dilution buffer, which was made using 5% (wt/vol) nonfat dried milk and 0.05% Tween 20 in PBS. Following the overnight incubation in primary antibody, plaque assays were washed with PBS three times and then incubated for 1 h in anti-mouse IgG horseradish peroxidase (HRP)-conjugated sheep antibody (GE Healthcare) diluted 1:4,000 in antibody dilution buffer. Assays were then washed three additional times with PBS and incubated in 0.5 ml of True Blue reagent for 30 min to allow for the staining of plaques. Once plaques were visible, plates were washed with water and allowed to dry before counting (only wells with greater than 3 plaques were used for the calculation of endpoint titer).

### Viral Growth Curves

For each growth curve, 200 PFU of respective virus was injected into 10-day old embryonated chicken eggs. Eggs were refrigerated at 0, 24, 48, or 72 hours post-infection and allatonic fluid was collected after 48 hours of refrigeration at 4°C. Aliquots of allantoic fluid were immediately frozen at −80°C to be thawed for use in HA assays and standard plaque assay procedures. All experiments were conducted in biological triplicate.

### Hemagglutination (HA) assay

Hemagglutination (HA) assays were performed by diluting virus-containing allantoic fluid in cold PBS. 50 microliters of chicken blood diluted 1:80 in cold PBS was mixed with each sample and incubated at 4°C overnight prior to scoring.

### DI unit calculation

Defective Interfering or “DI” Units were calculated by normalizing a virus’s HA score and endpoint titer to that of WT PR8. These normalized values were then averaged, and the HA score was divided by its normalized, averaged endpoint titer.

### Microscopy time course

12-wells of MDCK cells were seeded with approximately 85,000 cells for use 24 hours later for all microscopy experiments. MDCKs were infected for 1 hour at an MOI of 0.05 with either 9s PB1 mCherry, 9s PB2 sfGFP, or WT PR8 virus diluted in PBS/BSA at a total volume of 500 microliters. MDCKs were infected at an MOI of 0.1 with the 10s virus. The WT PR8 control was infected at an MOI of 0.05. Following the incubation period, the infection medium was removed and cells were placed in complete medium supplemented with 1:1000 diluted TPCK trypsin. At the indicated time after infection, MDCK cell medium was removed and replaced with 1 ml of warm PBS. Cells were incubated with Hoechst stain (1 microliter/ml of PBS) to allow for the staining of nuclei, and imaging was performed on the Zoe fluorescent cell imager (Bio-Rad) using the same gain, exposure and zoom settings for all images taken. Images were then processed with ImageJ (NIH).

## Supporting Information Legends

Supplemental Figure 1. Diagrams of the artificial viral segments tested in this study.

(A) Design of PB1 ORF flanked by NA packaging signals. (B) Design of mCherry flanked by PB1 packaging signals. (C) Design of PB2 ORF flanked by NP packaging signals. (D) Design of sfGFP flanked by PB2 packaging signals. (E) Design of the HA ORF flanked by NS packaging signals. (F) Design of sfGFP flanked by HA packaging signals. (G) Design of PB2 ORF flanked by PA packaging signals. (H) Design of M1 ORF flanked by HA packaging signals. (I) Design of the zsGreen (splice site) M2 ORF flanked by M packaging signals. (J) Design of the NS1 ORF flanked by NA packaging signals. (K) Design of the mCherry (splice site) NEP ORF flanked by NS packaging signals. For all diagrams, the indicated regions define the number of nucleotides. Dark grey regions represent silently mutagenized regions of the viral ORF.

