## Supplemental Figure for "Replication competent, 10-segmented influenza viruses as antiviral therapeutics"

**Supplemental Material: Haas, Harding and Heaton**

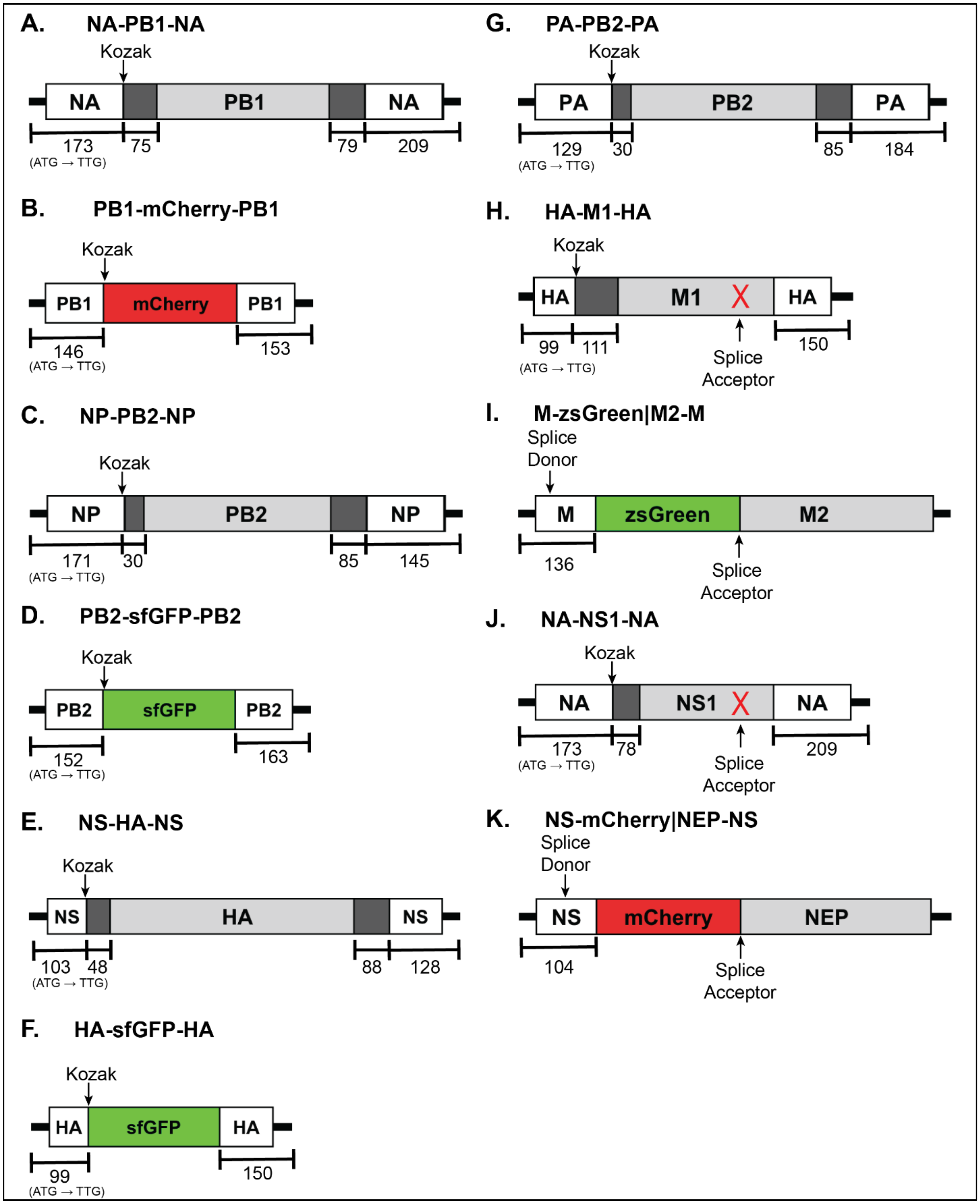


**Supplemental Figure 1. Diagrams of the artificial viral segments tested in this study.** (A) Design of PB1 ORF flanked by NA packaging signals. (B) Design of mCherry flanked by PB1 packaging signals. (C) Design of PB2 ORF flanked by NP packaging signals. (D) Design of sfGFP flanked by PB2 packaging signals. (E) Design of the HA ORF flanked by NS packaging signals. (F) Design of sfGFP flanked by HA packaging signals. (G) Design of PB2 ORF flanked by PA packaging signals. (H) Design of M1 ORF flanked by HA packaging signals. (I) Design of the zsGreen (splice site) M2 ORF flanked by M packaging signals. (J) Design of the NS1 ORF flanked by NA packaging signals. (K) Design of the mCherry (splice site) NEP ORF flanked by NS packaging signals. For all diagrams, the indicated regions define the number of nucleotides. Dark grey regions represent silently mutagenized regions of the viral ORF.
